# Current models cannot account for V1’s specialisation for binocular natural image statistics

**DOI:** 10.1101/497008

**Authors:** Sid Henriksen, Daniel A. Butts, Jenny C.A. Read, Bruce G. Cumming

## Abstract

A long-standing observation about primary visual cortex (V1) is that the stimulus selectivity of neurons can be well explained with a cascade of linear computations followed by a nonlinear rectification stage. This framework remains highly influential in systems neuroscience and has also inspired recent efforts in artificial intelligence. The success of these models include describing the disparity-selectivity of binocular neurons in V1. Some aspects of real neuronal disparity responses are hard to explain with simple linear-nonlinear models, notably the attenuated response of real cells to “anticorrelated” stimuli which violate natural binocular image statistics. General linear-nonlinear models can account for this attenuation, but no one has yet tested whether they quantitatively match the response of real neurons. Here, we exhaustively test this framework using recently developed optimisation techniques. We show that many cells are very poorly characterised by even general linear-nonlinear models. Strikingly, the models can account for neuronal responses to unnatural anticorrelated stimuli as well as to most natural, correlated stimuli. However, the models fail to capture the particularly strong response to binocularly correlated stimuli at the preferred disparity of the cell. Thus, V1 neurons perform an amplification of responses to correlated stimuli which cannot be accounted for by a linear-nonlinear cascade. The implication is that even simple stimulus selectivity in V1 requires more complex computations than previously envisaged.

**Significance statement:** A long-standing question in sensory systems neuroscience is whether the computations performed by neurons in primary visual cortex can be described by repeated elements of linear-nonlinear units (a linear filtering/pooling stage followed by a subsequent output nonlinearity, such as a squaring). This question goes back to the Nobel-prize winning work by Hubel & Wiesel who argued that orientation selectivity in V1 can qualitatively be explained in this way. In this paper, we show that V1 neurons have an amplification of their response to stimuli which are contrast matched in the two eyes, and that the recovered models cannot describe this property. We argue that this likely represents more sophisticated computations than can be compactly described by the linear-nonlinear cascade framework.

## Introduction

In their classic model for orientation selectivity in the visual cortex, Hubel and Wiesel (1962) proposed that simple cells performed a linear operation on the retinal image. A nonlinear relationship between the filter response and the spike rate was sufficient to account for the observed selectivity (a linear-nonlinear, or LN, model of simple cells [Hubel and Wiesel, 1962]). Complex cell properties such as position invariance could be modelled as the sum of several such LN “subunits” followed by a second nonlinearity. This process, where the output of one set of LN models forms the input to a second LN model, is referred to as an LN cascade. In the ensuing 50 years, models with this structure have been used to explain the responses of sensory neurons in many cortical areas, and increasingly sophisticated methods have evolved to fit the models to individual neurons. Disparity selectivity in these neurons is typically explained with the binocular energy model (BEM), an LN cascade model in which the linear filters are binocular [Ohzawa et al., 1990].

One interesting property of these neurons has been hard to capture with the model: when presented with anticorrelated random dot patterns, neuronal responses are less strongly modulated by disparity than in correlated patterns [Cumming and Parker, 1997]. This anticorrelated attenuation is thought to represent the fact that these stimuli cannot arise in natural viewing and hence represent a specialisation for the statistics of natural binocular inputs [Haefner and Cumming, 2008, Henriksen et al., 2016b].

Generalised versions of the BEM (GBEM), which allow for arbitrary filters and nonlinearities, as well as suppressive subunits, can capture a diverse set of tuning curves, including the attenuated response to anticorrelated stimuli [Nieder and Wagner, 2000, Read et al., 2002, Tanabe and Cumming, 2008, Tanabe et al., 2011, Henriksen et al., 2016c, Goncalves and Welchman, 2017]. However, none of these studies demonstrate that real cells obtain their disparity tuning through a GBEM-type mechanism.

A sufficiently broad LN cascade can approximate any function [Hornik, 1991], so in theory a GBEM with enough subunits must be able to describe the disparity tuning of real neurons. However, an LN cascade is only an interpretable description of a mechanism if the number of filters is modest. In practice, the number of subunits is also limited by the amount of data, since the cross-validated performance of a complex model will generally be low when there is limited data. The question then is whether real neurons can be described by a GBEM with a modest number of subunits.

The most comprehensive effort to answer this was by Tanabe et al (2011). They performed a spike-triggered analysis of covariance, using a binocularly uncorrelated Gaussian noise stimulus. GBEMs fitted to these data did not produce disparity tuning curves showing anticorrelated attenuation. However, this reflected the small correlation range present in the noise stimulus. When Tanabe et al separated their neuronal responses according to whether the stimulus correlation fluctuated above or below the mean of zero, they found that the neurons also showed no anticorrelated attenuation. It is then hardly surprising that the fitted models did not.

In this paper, we definitively answer the question of whether the GBEM framework can account for the disparity tuning of real cells. Using recently developed optimisation routines [McFarland et al., 2013], we fit GBEM units to neuronal data recorded from V1 in the macaque. The key advance from Tanabe et al (2011) is that this method allows us to fit GBEMs when the image sequences are binocularly correlated (or anticorrelated), and hence produce anticorrelated attenuation in the data. The GBEM accounted well for neuronal responses to anticorrelated stimuli, but underestimated response to correlated stimuli at the preferred disparity. This suggests that V1 contains a mechanism to amplify responses to naturally occurring disparities, which cannot be explained by an LN cascade.

## Materials and methods

### Animal subjects

Two male macaques (Macaca mulatta) were implanted with scleral search coils, head posts, and a recording chamber under general anaesthesia. The full procedure is described elsewhere [Cumming and Parker, 1999, Read and Cumming, 2003]. For the experiment, subjects viewed two CRT monitors through a custom mirror haploscope. The subjects were required to maintain fixation on a central box in order to receive a reward. All experiments were performed at the National Institutes of Health in the US, and complied with the US Public Health Service policy on the use and care of animals. The protocols received approval by the National Eye Institute Animal Care and Use committee at the National Institutes of Health.

### Recording

We recorded extracellular activity from neurons in primary visual cortex (V1) using laminar multi-contact electrodes (U-probes, Plexon for monkey Jbe; V-probes, Plexon for monkey Lem). Each electrode had 24 linearly arranged probes spaced 50 *μ*m apart. Electrodes were placed transdurally at the start of each day with a custom-built microdrive. The data was sampled with Spike2 (Cambridge Electronic Design), and the full waveform data was saved to disk for offline analysis. Spikes were subsequently analysed offline using custom spike-sorting software. Neurons were included in the analysis if they were well-isolated and disparity-tuned (as determined by a permutation test, *P* < 0.01). 95/197 cells met these criteria, giving 65 cells from monkey Lem, and 30 from monkey Jbe.

### Stimulus

The stimuli were presented on two Viewsonic P225f CRT monitors. The resolution was 1280 × 1024 pixels and the monitor refresh rate was 100Hz. Both monitors' luminance outputs were linearised using lookup tables. The mean luminance was 40 cd/m^2^, and the contrast (between maximum and minimum luminance) was > 99% as measured with a Konica-Minolta LS100 photometer. The eccentricity depended on the recording site, and varied from 2° – 4° for recordings from the operculum to > 10° for recordings in the calcarine sulcus. The animal was shown a 1D random noise pattern consisting of either black, gray, or white bars. The orientation of the stimulus was chosen so as to match the orientation preference of the cell (measured using circular patches of 1D noise at zero disparity) as closely as possible. When the stimulus orientation was sufficiently different from the preferred orientation (e.g. because there were multiple cells with different orientation preferences), there was typically no disparity selectivity to the 1D noise stimulus. The bars were 0.0946° in width, and the pattern consisted of 42 bars in each eye. The stimulus could be either binocularly correlated, anticorrelated, or uncorrelated. Stimulus disparities were selected based on the disparity tuning observed in measurements after fixing the orientation. Disparity was applied to the stimulus by wrap-around (i.e. bars displaced off the right end of the stimulus would be appended to the beginning of the stimulus; this has the effect of keeping the frequency power spectrum the same for the left and right images). We only used disparities which were integer multiples of the bar width, and always applied disparity orthogonally to the orientation of the bar pattern. A new stimulus pattern with a new disparity and correlation was shown every 30ms. A single trial lasted for 3s, corresponding to 100 independent noise patterns. We also implemented a two-pass procedure by duplicating trials, such that the same exact sequence of noise patterns occurred twice for most trials. This was done for the purposes of another experiment. Since there is no straightforward way of determining what was on a cell's receptive field prior to the onset of a trial, we discarded the first 200ms of each trial.

### Disparity tuning curves

In order to compute disparity tuning curves, we performed a forward correlation analysis. For a given disparity and correlation, we first identified all the patterns which were presented with the given stimulus parameters. For each pattern, we computed the number of spikes observed in bins *t*_max_ – 1, *t*_max_, and *t*_max_ + 1, where *t*_max_ is the time bin where we observed the largest variance across disparities. In other words, we computed the response in a 30ms window around the peak response for each neuron. The mean spike count was then computed for every disparity/correlation, giving a mean spike count for each disparity-correlation combination. The same exact procedure was performed for both the real cells and their model counterparts, with the only difference being whether the sequence of spike counts was predicted or observed.

### Generalised binocular energy model

We fit a generalised form of the binocular energy model using the framework developed by McFarland & Butts (2013). The model takes the general form

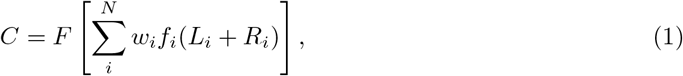

where *f_i_* is the subunit nonlinearity for the *i*^th^ subunit, *w_i_* is the weight given to the *i*^th^ subunit (constrained to be either −1 or +1, corresponding to a suppressive or excitatory subunit, respectively), and *F* is the final spiking nonlinearity of the model unit. *L_i_* and *R_i_* are the response of the *i*^th^ left and right filters, and are further defined as

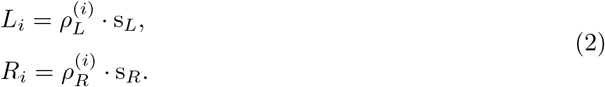

Here *s_L_* and *s_R_* refer to a vector representation of the stimulus presented to the left and right eyes, respectively. 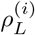 and 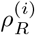 refer to vector representations of the linear spatiotemporal filters for the left and right eye of the *i*^th^ subunit. The number of spatial elements in the filter was simply the number of independent pixels in the stimulus, which was 42 for the left and right eyes (84 total). The number of temporal elements in the filter was 15, sampled at 10ms, corresponding to a maximum temporal kernel of 150ms. Thus, the total number of elements for the binocular spatiotemporal filter was 1260. The subunit nonlinearity *f_i_* can in principle take on a range of forms, but for the current purposes we have constrained it to always be a thresholded square. In symbols,

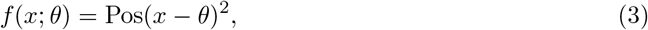

Where Pos refers to half-wave rectification, and *θ* is the threshold parameter. The spiking non-linearity *F* is a softplus rectifier, which is a smoothly varying rectifier function (with well-defined derivatives everywhere). It takes the form

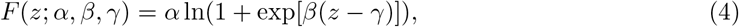

where *α, β*, and *γ* approximately correspond to magnitude, slope, and threshold parameters, respectively. The term in equation 1 is a continuous firing rate, and the spike count is obtained by passing the continuous rate through some discretisation procedure. For all analyses, we passed the continuous rate through a Poisson random number generator (poissrnd in Matlab) to obtain discrete spike counts.

### Hyperparameters

The optimisation routine developed by McFarland & Butts (2013) allows for fitting various components of this framework to empirical data by optimising the (log) likelihood of the model parameters given the data. Specifically, we can directly fit the coefficients of the linear spatiotemporal filters *L_i_* and *R_i_*, the thresholds *θ_i_* of the subunit nonlinearities, and the parameters *α, β*, and *γ* of the spiking nonlinearity. While the log likelihood surface is not guaranteed to be convex, McFarland & Butts note that appropriate steps, such as L1 and smoothness regularisation, can in general prevent the routine from converging to local minima [McFarland et al., 2013]. Therefore, we used both L1 and smoothness regularisation. L1 regularisation penalises the L1 norm (the taxicab distance from the origin) of the linear filter coefficients, ensuring that the filter coefficients take on sensible values and also encouraging sparsity of the filter coefficients [Tibshirani, 1996]. Smoothness regularisation penalises the Laplacian of the filter coefficients, ensuring that the second derivatives of the filters are small everywhere. This prevents abrupt, physiologically implausible changes in the filters (since such changes would correspond to large second derivatives). More generally, both forms of regularisation help prevent overfitting to the training data. The numbers of excitatory and suppressive subunits have to be optimised through cross-validation. In order to do this, we first split the data into a training set and a validation set. In order to prevent leakage of the training data into the validation data, the sets were split by trial instead of frame. For each cell, 75% of trials belonged to the training data, and the remaining 25% of trials belonged to the validation data. All identical trials (i.e. two-pass trials) were kept in the same set (either validation or training) to ensure independence of the two sets. We then performed a grid search on the number of excitatory and suppressive subunits, computing the log likelihood of the model on the validation data (the cross-validated log likelihood) for each combination of excitatory and suppressive subunits. Our hyperparameter search space was from 1 to 12 excitatory subunits, and from 0 to 5 suppressive subunits, yielding a total of 72 hyperparameter combinations for each cell. We capped the number of excitatory and suppressive subunits at 12 and 5, respectively, since we observed no cases of cells which were best modelled by 12 excitatory or 5 suppressive subunits. We repeated this procedure with different train/test splits at least 3 times for each excitatory-suppressive combination, and used the mean cross-validated log likelihood for model selection. The hyperparameter combination with the highest mean cross-validated log likelihood was used in the subsequent analysis. In general, the cross-validated log likelihood is fairly stable across iterations and so the best parameter combination is not greatly affected by either number of repeats or aggregation rule (i.e. mean, min, max, and median all give very similar results).

### Spike count predictions

After the hyperparameters of the model have been fixed, we obtained cross-validated model spike count predictions for each cell by running five-fold cross-validation. We first split the data into five equal subsamples (by trial, as noted above), and then fitted the GBEM on four of the subsamples. This allows us to obtain spike count predictions for hitherto unseen data (i.e. cross-validated predictions). We did this for each of the subsamples, resulting in cross-validated predictions for all trials in our dataset. This ensures that all model predictions are of unseen data. Effectively, we are testing a particular model architecture (i.e. the GBEM with a certain number of excitatory and suppressive subunits) as opposed to a particular instantiation of the GBEM.

### Model responses to disparity-correlation combinations

The disparity tuning curve captures how well a GBEM unit can capture responses across disparities. In order to describe how the GBEM units captures the cell’s responses *within* disparities, we first identify frames with the appropriate disparity/correlation, and then run the forward correlation procedure as previously specified. The same exact procedure was performed for observed spike counts as for (cross-validated) predicted spike counts.

Suppose the experiment includes N frames with a given disparity and correlation, where we typically have *N* > 1000. The forward correlation analysis gives us *N* predicted spike counts obtained from the set of cross-validated models. The average of these spike counts gives us the predicted disparity tuning curve. However, both cells and models show considerable variability in the response to different stimuli with the same disparity and correlation. To assess the agreement *within* disparities, we first binned the *N* predicted spike counts into 15 bins chosen so as to contain an equal number *N*/15 of spike counts. This allows us to get both the mean model spike count within each bin, and, using the same bin boundaries, the mean spike count using the counts observed for the real neuron. If we plot the model spike count against the cell spike count in this way, we would expect the points from an unbiased model to lie on the identity line.

### Binocular Gaussian noise

In order to compare our model responses to the population responses of cells in Tanabe & Cumming (2011), we also computed the responses of the model units to binocular Gaussian 1D noise. Each pixel in these stimuli had a value which was drawn from a normal distribution with unit variance, and was independent in the two eyes and also independent from frame to frame. A new pattern was generated every 10ms. In order to construct “tuning curves” for the independent binocular Gaussian noise, we used a procedure like that used by Tanabe et al (2011). We first computed the normalised binocular cross-correlation function of each image frame, extracting a correlation value for each disparity. For each disparity, we then identified the frames with the top and bottom 20% of correlation values, which correspond to our correlated and anticorrelated frames, respectively. We then used these frames to trigger the forward correlation procedure, computing disparity tuning curves as previously specified. With this procedure, a single frame can be used in multiple disparities if the magnitude of binocular correlation exceeded our threshold for more than one disparity.

### Experimental design and statistical analysis

We used a permutation test to assess the significance of disparity tuning in a given cell. For comparing the GBEM's ability to capture the cell's response to different stimuli, we first compute the relevant metrics (e.g. correlated and anticorrelated tuning strength, anticorrelated attenuation, as defined in the Results), and then do a paired samples t-test. We also carried out Pearson correlations, obtaining both *r* and *p* values for the relationship between these metrics. In Figure 2, bootstrap confidence intervals (CIs) are computed on the spiking response of both the cell and the model. The code for permutation tests and bootstrap CIs was written by the authors in MATLAB, while the Pearson correlation and the paired samples t-tests were carried out using MATLAB’s `corr` and `ttest` functions, respectively.

**Figure 2:**
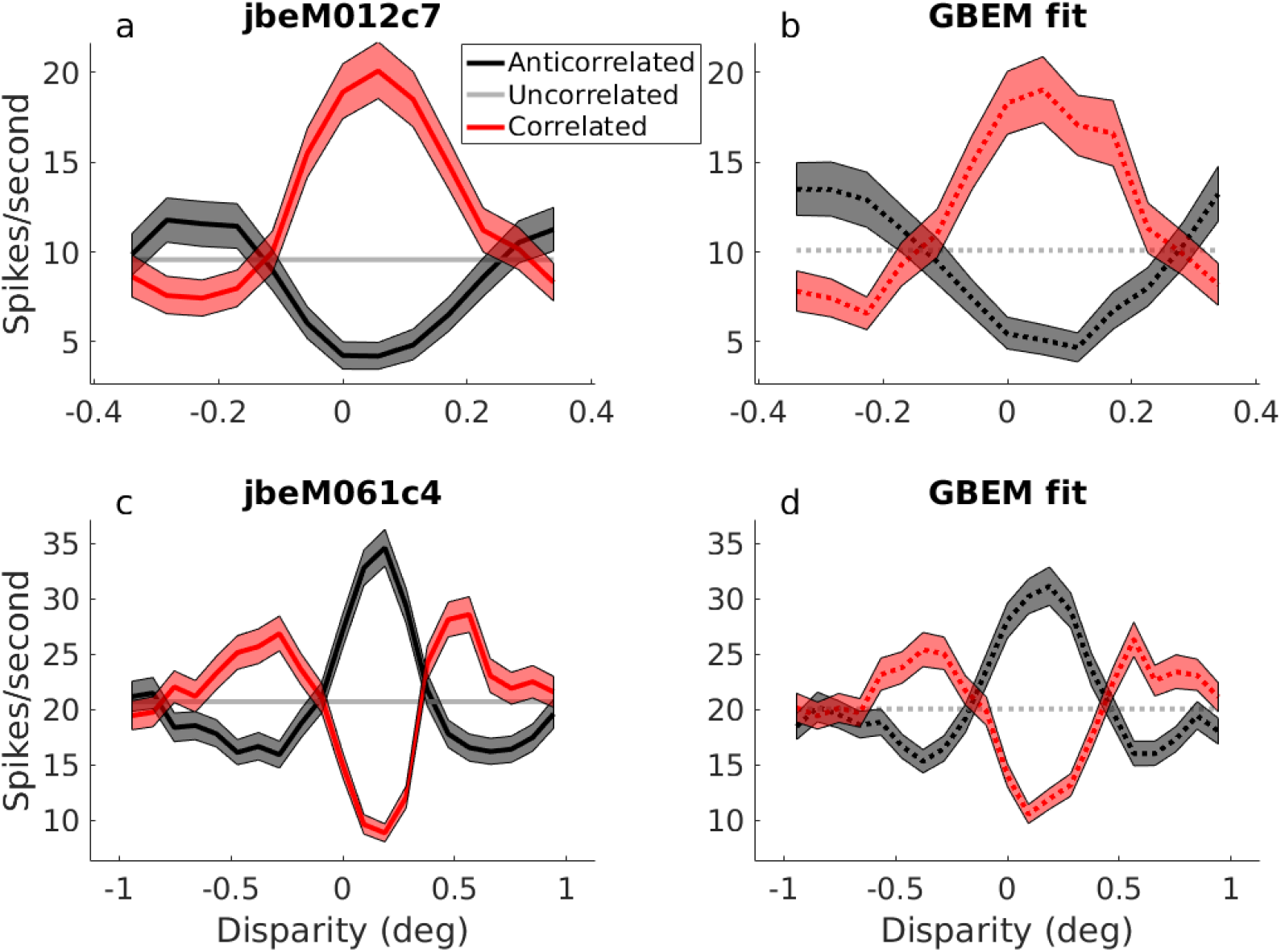
Disparity tuning curves for two example cells (a,c) and their corresponding GBEM fit (b,d). Correlated responses are shown in red, anticorrelated responses are shown in black, and uncorrelated responses are shown in gray. The number of spikes per second were calculated in a 30ms window centred around the peak temporal response of the cell. The shaded regions show 95% bootstrap confidence intervals for the responses.

## Results

### Example LN filters

From the fitting procedure, we obtain a GBEM unit with an optimal number of excitatory and suppressive subunits for each cell. The linear filters for a GBEM fit to cell lemM322c1 are shown in Figure 1a. For this cell, all subunits have a phase disparity between the two eyes (i.e. the profile of the filters differ in the left and right eyes). Figure 1b shows the subunit nonlinearities for the excitatory and suppressive subunits. In this case, the fitted thresholds are similar for the different subunits, except for one of the excitatory subunits (bronze line in top panel of Figure 1b). While this subunit's filter coefficients are much lower (excitatory subunit with bronze outline in Figure 1a), its threshold is much more negative, meaning that this subunit gives a large positive output to a blank screen. Figure 1c shows “tuning curves” to correlated stimuli for the excitatory pool and the suppressive pool (i.e. summed over excitatory and suppressive subunits, respectively). The pooled responses are normalised such that the baseline (median) response is zero (necessary for visualisation purposes since the excitatory subunit in Figure 1b has a high baseline response). The tuning curves of the excitatory and suppressive pools have opposite phases: this is the familiar push-pull organisation for binocular neurons introduced by Read & Cumming (2007) and found in V1 neurons by Tanabe, Haefner, & Cumming (2011).

**Figure 1:**
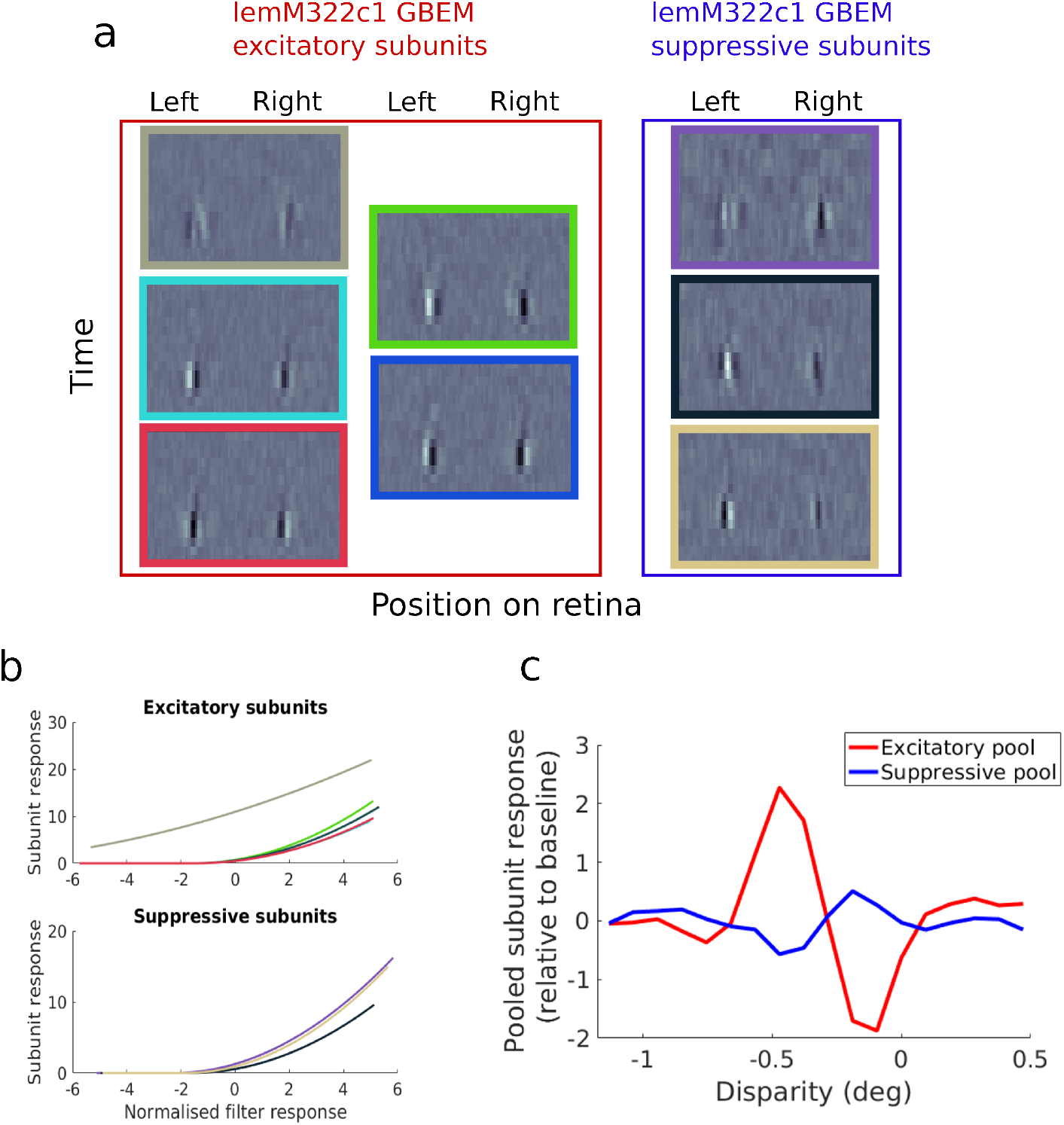
a) Filters of excitatory and suppressive subunits recovered for cell lemM322c1. The resulting GBEM unit has 5 excitatory subunits (red outline), and 3 suppressive subunits (blue outline). The filters are shown here as spatiotemporal filters, with the vertical axis denoting time and horizontal axis denoting space. Note that half of the pixels correspond to the left eye and the other half to the right eye (separated by a vertical bisection). b) Subunit responses as a function of the normalised filter response for both excitatory (top) and suppressive (bottom) subunits. The filter response will necessarily be centred on zero. In order to normalise the filter responses, we divide by the standard deviation. The normalised filter response is therefore a z-score where a value of ±1 corresponds to one standard deviation from the mean (which as noted is necessarily zero). A blank screen corresponds to a filter response of 0 in this scheme (though the converse is not true: a filter response of 0 does not necessarily imply a blank screen). Each line shows a different subunit, with the colours mapping onto the outline of the spatiotemporal filters in a). c) Disparity tuning curves for the excitatory (red) and suppressive (blue) pools. The baseline (median) response of the pools has been normalised to zero so that a meaningful comparison can be made. The excitatory pool has a much higher baseline response than the suppressive pool.

### Example disparity tuning curves

Once we have the GBEM fits for each cell, we can obtain a disparity tuning curve for both the cell and the model, as described in the Methods. Figure 2 shows the disparity tuning curve for two cells where the GBEM unit has successfully captured the disparity tuning of the cell. Figure 2a shows the tuning curve for cell jbeM012c7, a “tuned excitatory” cell by Poggio & Fischer’s (1977) nomenclature. Figure 2b shows the corresponding GBEM fit. It is important to highlight that the model was not fit to the disparity tuning curve of the cell; rather, the optimisation routine was given the luminance values on the screen and the spike times. Furthermore, the model disparity tuning curves used only spike counts predicted for on-screen luminance values which were not used during fitting. Thus, the ability to capture disparity tuning means that the model has learned a nonlinear binocular interaction which was not explicated in the inputs. The GBEM does a good job at capturing both the correlated and anticorrelated responses of the cell, and also captures the weak response attenuation of the cell to anticorrelated stimuli. Figure 2c shows the tuning curve of another example cell of the type known as a “tuned inhibitory” cell [Poggio and Fischer, 1977]. These cells are relatively uncommon in cortex and respond most vigorously to a stimulus which is binocularly anticorrelated, i.e. to stimuli which are impossible in naturalistic viewing. A success of the original BEM is that it can capture this type of disparity tuning by incorporating phase disparity between the left and right subunits. Indeed, the GBEM readily accounts for the disparity tuning of this cell (Figure 2d), and does so by recovering linear filters which have phase disparities of approximately *π*.

Most GBEM fits capture the overall shape of disparity tuning, and for some cells the magnitude of disparity tuning is also well-captured (e.g. Figure 2). However, most GBEM units underestimate the magnitude of disparity tuning. This becomes particularly evident when the GBEM and cell responses are superimposed, as in Figure 3. These include example cells where the disparity amplitude is quite well captured (Figure 3a), significantly underesimated (Figure 3b, d) and very seriously underestimated (Figure 3c). Figure 3a shows an example odd-symmetric cell where the cell’s response is very well-captured.

**Figure 3:**
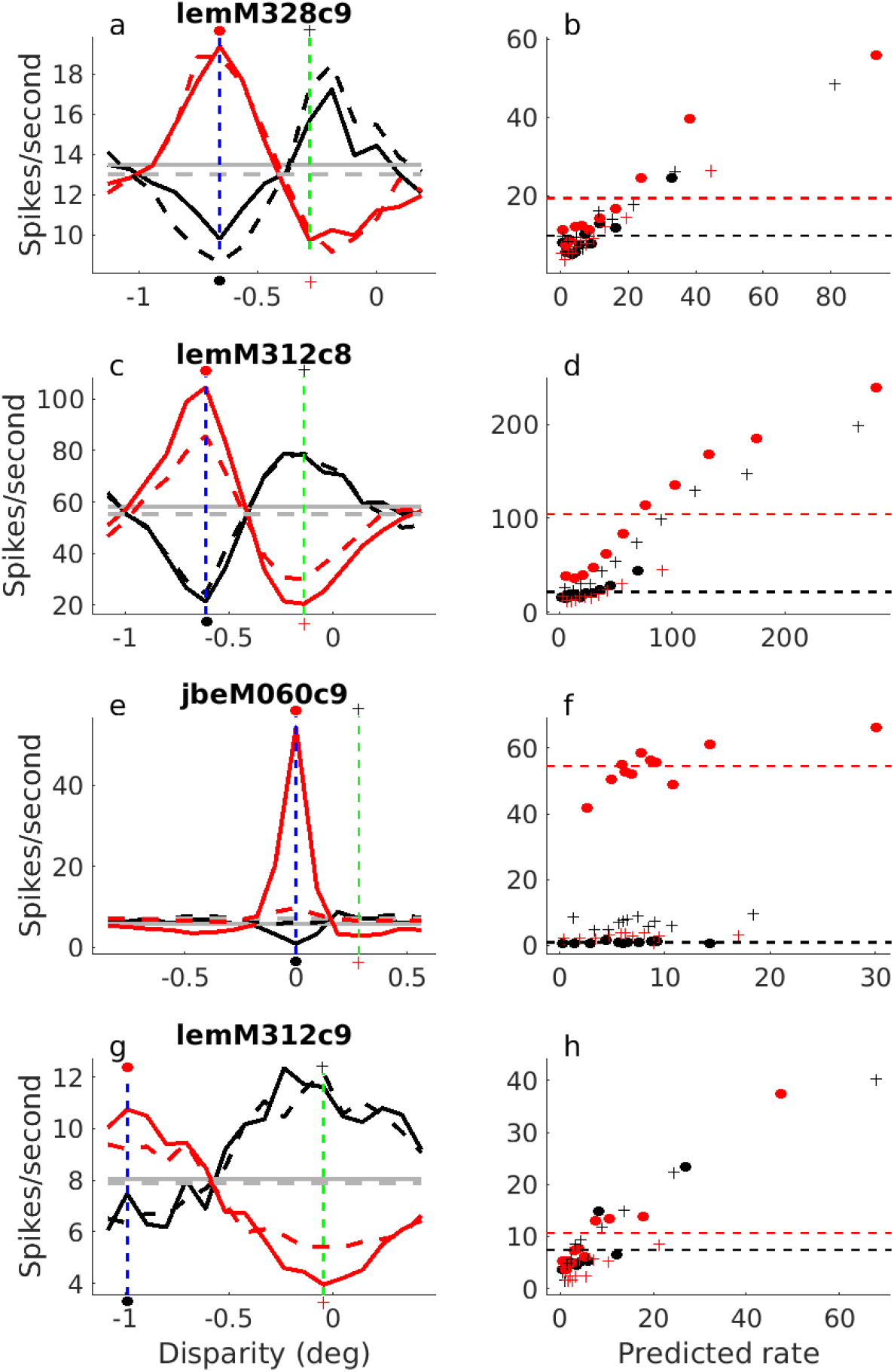
Disparity tuning curves for three example cells which capture the diversity of model fits. Here, the response of the model (dashed lines) is overlaid on the response of the cell (solid lines). Panels a, c, e, and g show the disparity tuning curves, whereas panels b, d, f, and h show the observed spike rated plotted as a function of the predicted spike rate (binned according to the predicted spike rate). Each point in b, d, f, and h shows the average observed and predicted rates for a separate bin for a particular disparity/correlation combination (black dots are anticorrelated, red dots are correlated). Filled circles show the preferred disparity of the cell (dashed blue line in the left column), and pluses show the anti-preferred disparity (dashed green line in the left column). The horizontal dashed lines show the mean spike counts at the preferred disparity, for correlated (red) and anti-correlated (black) stimuli. For clarity, black and red dots and crosses are shown on the tuning curves to highlight the where the binned responses in b, d, f, and h are computed. In other words, the red dots in b are the binned responses of the disparity/correlation indicated by the red dot in a (and the average spike/second for the red dots in b is equal to the red dot in a).

### In exampe cells, models tend to underestimate disparity tuning amplitude

The response shown for all model cells is cross-validated, meaning that the model responses are shown for stimuli with which the model has not been fit. Thus, despite the underestimates, Figure 3 represents a substantial success of the GBEM architecture to account for average responses across disparities.

However, as noted, the amplitude of disparity tuning is systematically underestimated, especially at the preferred disparity, indicated with the vertical blue dashed lines in Figure 3a, c, e, and f. In order to gain insight into this discrepancy, we plotted the mean rate in the real cell as a function of the predicted rate of the model, binned according to the predicted rate. This relationship is shown separately for 4 different conditions: correlated and anticorrelated stimuli at the preferred and anti-preferred disparity (see Methods). If the model correctly predicts the firing rate of the cell across all conditions, all points should lie on the identity line. If the model correctly predicts responses across noise patterns within a disparity, then these points should be monotonically increasing within each disparity.

In Figure 3a and 3g, the disparity tuning curves are generally well captured, with only a slight underestimate. Accordingly, in Figure 3b and 3h, the points lie close to the identity line, indicating a generally successful model. In Fig 3c, the model captures the overall shape of the tuning curve, but underpredicts the magnitude of disparity tuning. In particular, it underpredicts the response of the cell to correlated stimuli. The effect of this can be seen in Figure 3d, where the responses to the correlated anti-disparity (red crosses) and the anti-correlated preferred disparity (black dots) lie on roughly the same curve. The correlated preferred responses are notably shifted up and to the left, i.e. the real cell responds more strongly to the preferred disparity than predicted by the model. This means that with the recovered model structure, there is no single output nonlinearity that can simultaneously account for the responses to correlated and anticorrelated stimuli. The implication is that the GBEM structure itself is not appropriate for simultaneously describing the response to correlated and anticorrelated stimuli. It also suggests that the correlated responses at the preferred disparity are in some way “special” in the way they are processed by this V1 neuron.

In some cases, we observed a much more spectacular failure to capture the disparity tuning. Figure 3e shows a cell where the model exhibits very little disparity selectivity. Despite this failure to capture disparity tuning, Figure 3f shows that the rank ordering of the model responses within disparities are nevertheless broadly preserved: stimuli of a particular disparity which elicit a larger than average response in the cell also elicit a larger than average response in the model. In other words, the model can account for a large proportion of the cell’s responses to variations in the stimulus, but simply fails to account for disparity-selectivity. This means that for a given disparity, a large response in the cell to a particular luminance pattern does generally correspond to a larger response in the model as well. However, the GBEM selectively fails to capture the binocular interaction between the left and right eyes. We found that in general, the model fits can be broadly grouped into the three categories highlighted in Figure 3, namely: disparity tuning well captured; captured with signficnant underestimate of amplitude; not captured at all. Of particular interest are the model units that fail to capture any disparity tuning (e.g. Figure 3e); this does not represent a general fitting failure, as evidenced by Figure 3f, but is, as noted, a selective failure to capture disparity tuning. If a GBEM unit failed to capture less than 30% of the variance in disparity tuning of the corresponding cell, we categorised this model fit as failing to adequately capture disparity tuning. Out of the 95 disparity-selective cells in our dataset, 28 fits failed to capture disparity tuning by this definition (30% of disparity-selective cells).

This failure is surprising given that, as noted above, GBEM units can easily produce disparity tuning curves of the required form. Indeed, it would be easy to hand-tune a GBEM to produce a much better fit to the tuning curve in Figure 3e than produced by our fitting procedure. It might therefore appear that this represents a trivial failure of our fitting procedure, e.g. convergence on a local optimum rather than the global one for a particular combination of excitatory and suppressive subunits. This is not the case. The reason for the “failure” to capture disparity tuning is that the model is not fit to the disparity tuning curves. Instead, the input to the model is simply the luminance patterns on the screen, and so the GBEM attempts to capture the full range of monocular, binocular, and temporal dynamics of the cell. The response to a given disparity in a tuning curve averages across many different monocular stimuli. Modest failures to capture responses to indiviudal stimuli can therefore result in substantial failures to describe the disparity tuning curve. Thus, while a hand-tuned model might be able to capture the tuning curve, it would do much worse than the fitted GBEM in predicting the response of the cell to actual (previously unseen) random line stereograms.

### Quantifying the reduced response to anticorrelated stereograms: the relative anticorrelated response

The total failure to capture disparity tuning, seen in about 30% of our cells, can be viewed as an extreme version of a more general failure: the underestimation by the GBEM of the magnitude of disparity tuning in real cells. Thus, the largely absent disparity tuning of the GBEM unit for jbeM060c9 (Figure 3e) can be seen as a more extreme failure of the sort seen in lemM312c8 (Figure 3c). Further evidence that the correlated response is special can be seen in the tuned inhibitory cell shown in Figure 3g. This cell’s response is suppressed by stimuli at disparity near 0, yet the GBEM unit underpredicts the magnitude of response modulation to these stimuli. This suggests that the model fails to capture the cell’s response to correlated stimuli *per se* and not just the spike rate independent of stimulus characteristics. The anticorrelated attenuation depends on the magnitude of the correlated response, which is amplified in real cells. Therefore, the effect of underestimating the response to correlated stimuli (at the preferred disparity) is that real cells have stronger anticorrelated attentuation than the GBEM units.

In order to quantify this, we compute the regression slope (type II regression, [Draper and Smith, 2014]) between the correlated and anticorrelated response. We will refer to this metric as the relative anticorrelated response of the cell (note that this is different from the metric used to quantify anticorrelated attenuation in [Cumming and Parker, 1997], but the same as that used by [Henriksen et al., 2016b]). Figure 4b shows graphically how the relative anticorrelated response quantifies the degree to which dispartiy tuning is reduced for anticorrealted stimuli. If the relative anticorrelated response is −1, then the neuron responds as strongly to anticorrelated stimuli as to correlated, but with a sign inversion (as in the standard binocular energy model) and thus exhibits no response attenuation to anticorrelated stimuli. A relative anticorrelated response of 0 corresponds to a neuron which does not modulate its response to anticorrelated stimuli ^1^.

**Figure 4:**
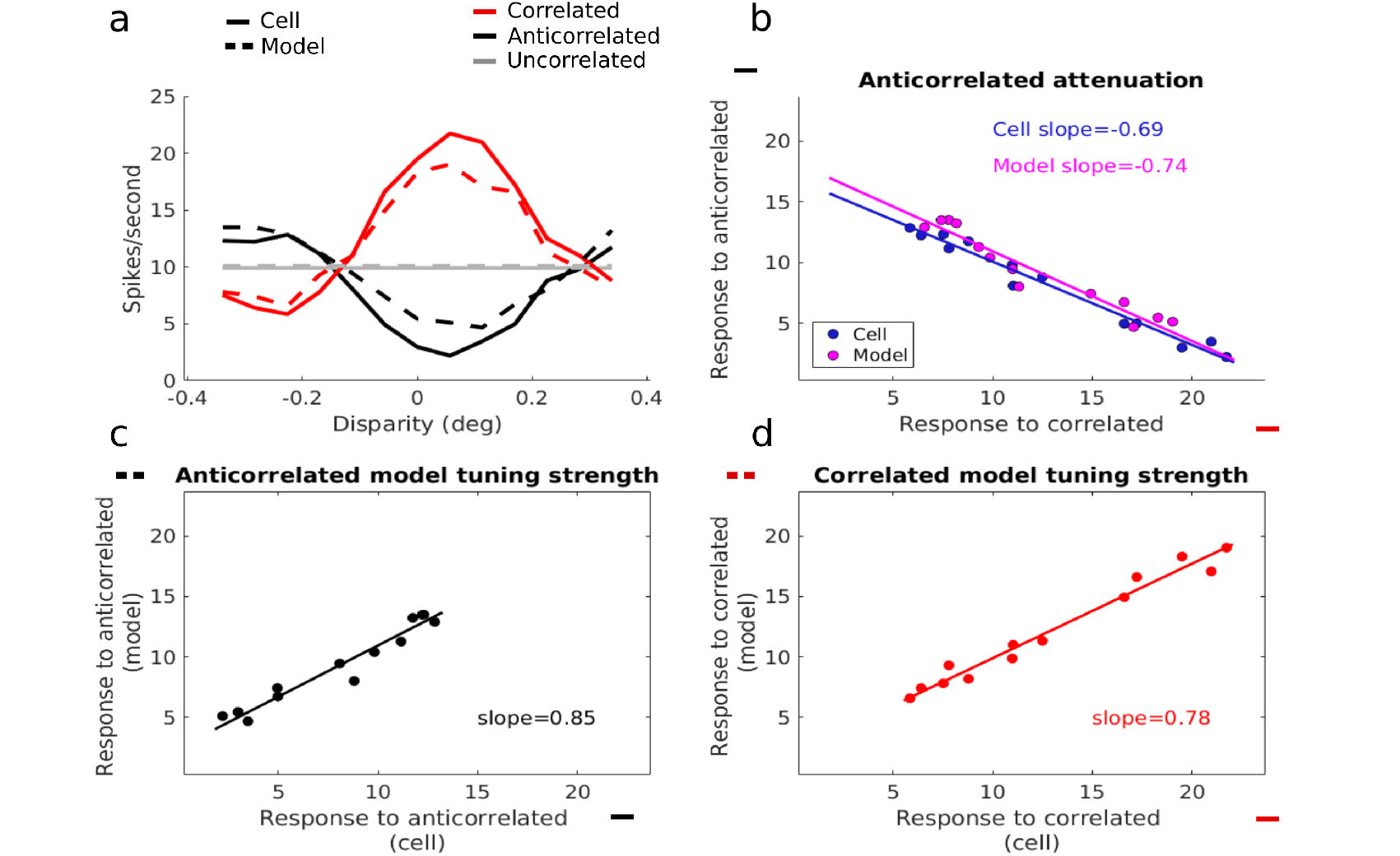
a) A disparity tuning curve for jbeM012c7 (solid lines), and its model fit (dashed line). b) Anticorrelated response as a function of the correlated response for a cell (blue dots) and its corresponding GBEM fit (magenta dots). The relative anticorrelated response is defined as the slope of the (type II) regression line, shown for both the model and cell. A relative anticorrelated response of 0 means that the cell does not modulate its response to anticorrelated stimuli, whereas an relative anticorrelated response of −1 means that the cell inverts its response perfectly just like in the binocular energy model. c) Anticorrelated response of the model as a function of the anticorrelated response of the cell. The regression slope defines the anticorrelated model tuning strength, which quantifies how strongly the model unit is tuned to anticorrelated stimuli relative to the cell. If the anticorrelated model tuning strength is 0.5, then the model unit modulates its response half as strongly to disparity for anticorrelated stimuli compared to that of the cell. d) Correlated response of the model as a function of the correlated response of the cell. The regression slope defines the correlated model tuning strength, which quantifies how well the model’s shape and magnitude of disparity tuning matches that of the cell.

The relative anticorrelated response for jbeM012c7 (Figure 2a) is −0.69 (95% CI: [−0.76, −0.62]), and the relative anticorrelated response for its GBEM fit (Figure 2b) is −0.74 (95% CI: [−0.85, −0.63]). Thus, the GBEM readily captures much of the anticorrelated response of this cell. For jbeM061c4 (Figure 2c), the GBEM also captures the relative anticorrelated response of the neuron well (Figure 2d), and this is again reflected in the relative anticorrelated response (jbeM061c4: −1.01, 95% CI: [−1.09,−0.923]; GBEM: −1.11, 95% CI: [−1.21, −1.00]). For the example cells in Figure 3, the relative anticorrelated responses are −0.57 [−0.75,−0.2] for lemM328c9 and −0.94 [−1.02,−0.86] for its GBEM fit; −0.67 [−0.71,−0.63] for lemM312c8 and −0.99 [−1.05,−0.93] for its GBEM fit; −0.13 [−0.14,−0.12] for jbeM060c9 and −0.62 [−0.70,−0.54] for its GBEM fit; −0.88 [−1.08,−0.70] for jbeM012c9 and −1.25 [−1.46, −1.07] for its GBEM fit.

### The GBEM fails to capture the anticorrelated attenuation seen in real neurons

In order to summarise the inability of the model to capture the cell’s tuning curve across the population, we plot the relative anticorrelated response for the neurons against the relative anticorrelated response for their corresponding GBEM fits (Figure 5a). The four example cells in Figure 3 are indicated. Two key points are worth observing. First, there is a strong positive correlation between the relative anticorrelated response in the cells and that seen in the model units (*r* = 0.61,*p* < 0.001, Pearson correlation). This is notable since Tanabe & Cumming (2011) found no attenuation in anticorrelated response in the model units recovered with their spike-triggered analysis of covariance. Thus, although it has long been recognised that the GBEM can in principle explain anticorrelated attenuation, this is the first direct evidence that this explanation is at least partially correct. However, the vast majority of points in Figure 5a lie beneath the identity line, meaning that the GBEM systematically predicts less anticorrelated attenuation than is observed in the cells. This suggests that although the GBEM goes some way towards accounting for the relative anticorrelated response seen in V1 neurons, it is not on its own a sufficient explanation. Notably, most of the cells that are above the identity are the ones we previously identified as producing poor fits to the disparity tuning curve of the cell.

**Figure 5:**
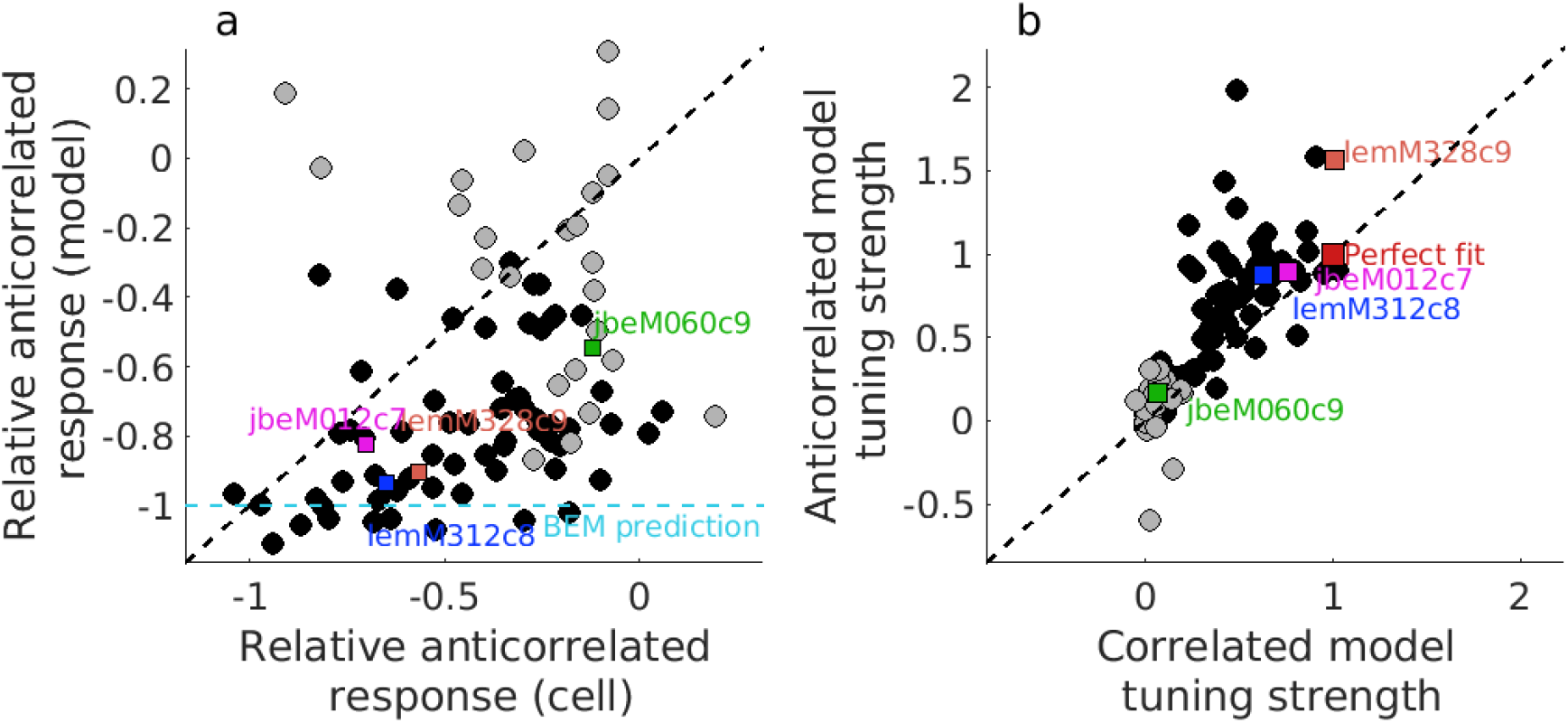
The relative anticorrelated response for the model against the relative anticorrelated response for the cell. The dashed black line shows the identity line, while the dashed cyan line shows the prediction of the BEM (perfect inverted tuning curve to anticorrelated). While the model shows some attenuation (values > −1), this is more dramatic in the neuronal data (the points lie mostly below the identity line). The model tuning strength for the anticorrelated tuning curve, against the same for the correlated tuning curve. The dashed black line again shows the identity line, and the red square marker shows the point of perfect model fit. Cells which produce poor GBEM fits (< 30% accounted for variance in the disparity tuning curve) are shown in gray in both plots.

### Quantifying model fit: the correlated and anticorrelated tuning strength

In principle, there are three potential reasons for the failure of the GBEM to account for anti-correlated attenuation. The first possibility is that the GBEM fails because it is unable to account for the anticorrelated response, perhaps because cells have a specialised mechanism for suppressing false matches [Samonds et al., 2013]. The second possibility is that the GBEM is unable to capture the correlated response. Similar to the anticorrelated case, a failure to capture the correlated response could be because V1 neurons have enhanced responses to true matches which cannot be easily captured by the GBEM framework. The third possibility is that there is failure to capture both the anticorrelated and the correlated responses. In our example cells above, we saw that the evidence pointed to the second possibility – a failure to capture correlated response at the preferred disparity. To quantify this at the population level, we first define a model tuning strength metric separately for the correlated and the anticorrelated patterns. The model tuning strength is simply the regression slope between the cell and the model’s tuning curves (illustrated in Figure 4c and d for the anticorrelated and correlated case, respectively). If the model tuning strength is 0.5, then the model’s disparity tuning is 50% of the real cell’s tuning (provided that the shape of the disparity tuning curve has been appropriately captured, which is generally the case in our data). We can compute the model tuning strength separately for correlated and anticorrelated responses, yielding a metric for how well a GBEM unit is able to capture the shape and magnitude of the cell’s disparity tuning for correlated and anticorrelated stimuli. Figure 5b shows the anti-correlated model tuning strength on the vertical axis, and the correlated model tuning strength on the horizontal axis. The vast majority of cells in this plot lie above the diagonal, suggesting that the GBEM is better able to capture the anticorrelated responses than the correlated ones. This was confirmed by a paired samples t-test comparing the anticorrelated model tuning strength (*M* = 0.52) against the correlated model tuning strength (*M* = 0.32): *t*(94) = 5.90, *P* < 10^−7^. The fact that points generally lie below 1 for both correlated and anticorrelated stimuli means that the models underpredict the strength of disparity tuning for both correlated and anticorrelated stimuli. However, the magnitude of the model failure is notably greater for correlated stimuli compared to anticorrelated stimuli.

The previous analysis confirms that the correlated responses of the cells are more problematic for the GBEM than anticorrelated responses. This is noteworthy since it suggests that the established way of thinking about the shortcomings of the BEM – that it is not able to capture the anticorrelated response – is incorrect. Instead, real neurons tend to have a larger response to true matches at one specific preferred disparity, which neither the BEM nor its generalisation (the GBEM) is able to capture. If the failures of the GBEM above reflect the effect of a mechanism that amplifies responses to correlated patterns, it predicts a sytematic relationship between this failure and relative anticorrelated response observed in the neuronal responses. Figure 6a shows the model tuning strength for correlated responses against the relative anticorrelated response in the cell. There is a strong negative correlation between the two (*r* = −0.58, *p* < 10^−9^, Pearson’s r). That is, the disparity tuning curves for correlated stimuli are systematically captured less well for cells which show stronger attenuation to anti-correlated stimuli. Figure 6b shows the equivalent plot for the model tuning strength to anticorrelated responses. This relationship is weaker, but significant (*r* = 0.34, *p* = 0.001). Interestingly, the effect in Figure 6b is largely due to cells such as jbeM060c9 which fail to account for the disparity tuning curve of the real cells. These are shown highlighted in gray and largely cluster in the bottom right of the plot. Excluding cells with bad fits (e.g. those that can account for less than 30% of the variance in the cell’s disparity tuning curve, gray circles in Figure 6b), makes the relationship non-significant (*r* = 0.19, *p* = 0.13). The results in Figure 6a remain highly significant (*p* < 10^−7^) when excluding poor fits.

**Figure 6:**
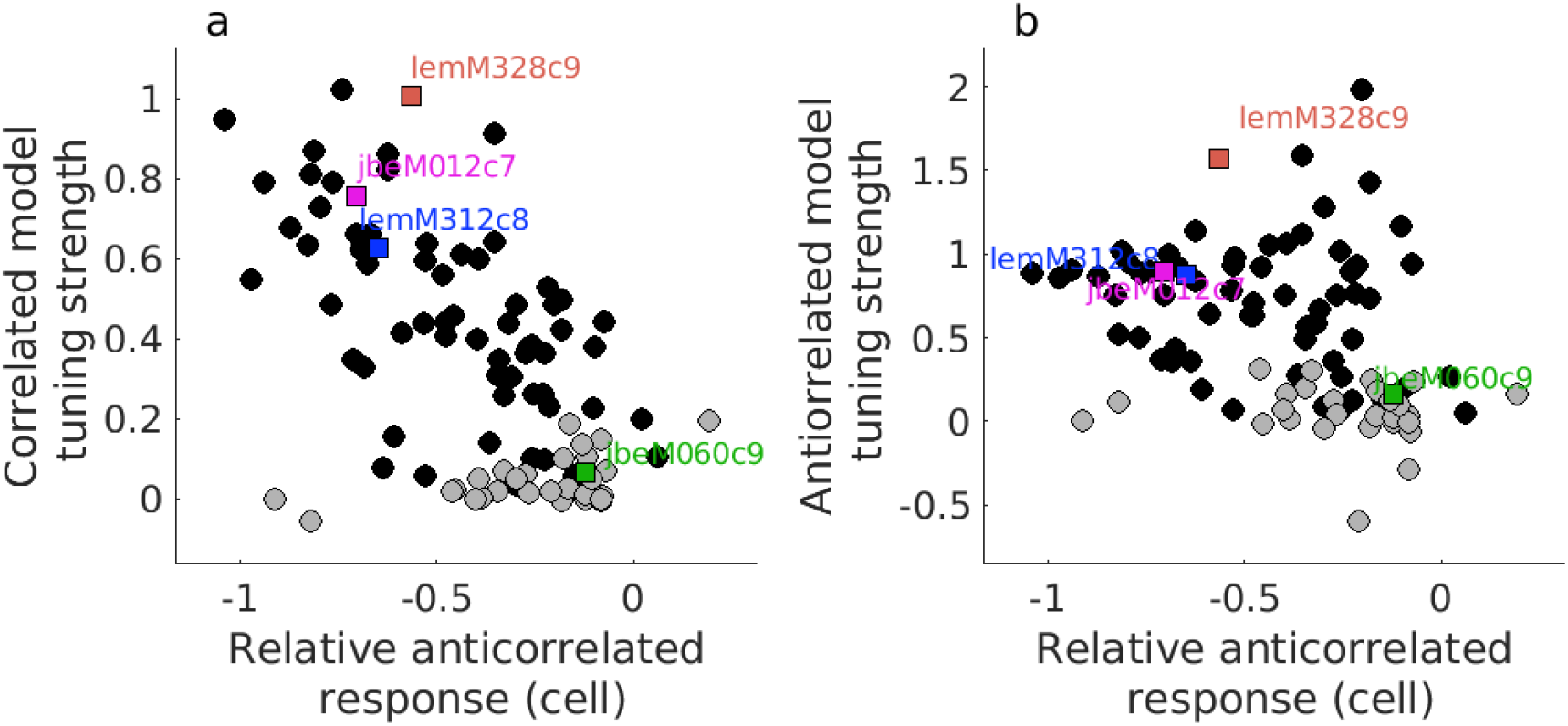
Correlated (a) and anticorrelated (b) tuning curve slope plotted as a function of the cell’s relative anticorrelated response. Tuning curve slope quantifies how well the magnitude of disparity tuning in the cell is captured by its GBEM fit, and this can be computed separately for the correlated and anticorrelated tuning curves. While the model’s correlated tuning strength is significantly related to the cell’s anticorrelated slope (a), the model’s anticorrelated tuning strength (b) is not. In other words, a cell’s magnitude of anticorrelated attenuation (i.e. its relative anticorrelated response) is predictive of whether the correlated response will be well-described by a GBEM unit. Cells which produce poor GBEM units (< 30% variance accounted for in the disparity tuning) are shown in gray.

Taking the results of Figure 5 and 6 together suggests that “attenuated anticorrelated response” is the wrong way to think about the response properties of disparity-selective cells. Instead of “attenuated anticorrelated response”, real neurons show “correlated amplification”: that is, they have an amplification of their response to true matches (correlated stimuli at the preferred disparity of cell) rather than an attenuation or suppression of false matches per se. This amplification is largely responsible for the failure of the GBEM to capture the disparity tuning curve of real cells.

### Model units correctly predict lack of anticorrelated atenuation with binocular Gaussian noise stimuli

The most comprehensive attempt at directly modelling disparity-selective cells in V1 by learning model parameters from data was performed by Tanabe & Cumming (2011). In order to retrieve the linear filters for each subunit, the authors first performed a spike-triggered analysis of covariance, and then optimised the nonlinearities for each subunit. In principle, this approach should give very similar results to the analysis we show here. It is noteworthy then that Tanabe & Cumming (2011) found no reduction in relative anticorrelated response in their model cells when tested on disparate 1D line patterns, where the real neurons exhibited strong reductions. The key differences between that paper and the current work are 1) the method of model estimation, 2) the duration of each individual noise pattern (10ms in Tanabe & Cumming, 2011; 30ms in the present work), and 3) different stimuli due to the requirements of their spike-triggered covariance method. Specifically, the spike-triggered analysis of covariance approach used by Tanabe & Cumming requires a white noise stimulus which is independent in the two eyes. The consequence of using independent noise in the two eyes is that there are very few frames with extreme binocular correlation values (e.g. close to −1 or 1). Interestingly, the authors showed that when the cells were tested on the same independent noise patterns with which the models were fit, the cells did not exhibit systematic attenuation to anticorrelation either [Tanabe et al., 2011].

An independent test of our GBEM units is then to explore whether they can reproduce this characteristic feature of real cells highlighted by Tanabe & Cumming (2011): real neurons show anticorrelated attenuation only to large positive or negative correlation values. The stimulus used to fit the GBEM is very similar to the RLS stimuli used by Tanabe & Cumming to construct their attenuated tuning curves, so we know that the model units show somewhat attenuated anticorrelated response (or more appropriately, correlated amplification) in this case. Thus, an additional test is whether our model units also show an absence of that attenuation to independent binocular Gaussian noise. The procedure for calculating disparity tuning curves from independent noise data was the same as in Tanabe & Cumming (2011) and is documented in the Methods section. Figure 7a shows a tuning curve computed using Tanabe & Cumming’s independent noise stimulus for the example GBEM unit shown in Figure 2a (jbeM012c7). While the model unit previously showed clear anticorrelated attenuation with disparate line stereograms (M=−0.74, 95% CI: [−0.85, −0.63]), this is now not significantly different from −1 (M=−0.95, 95% CI: [−1.08,−0.82]).

**Figure 7:**
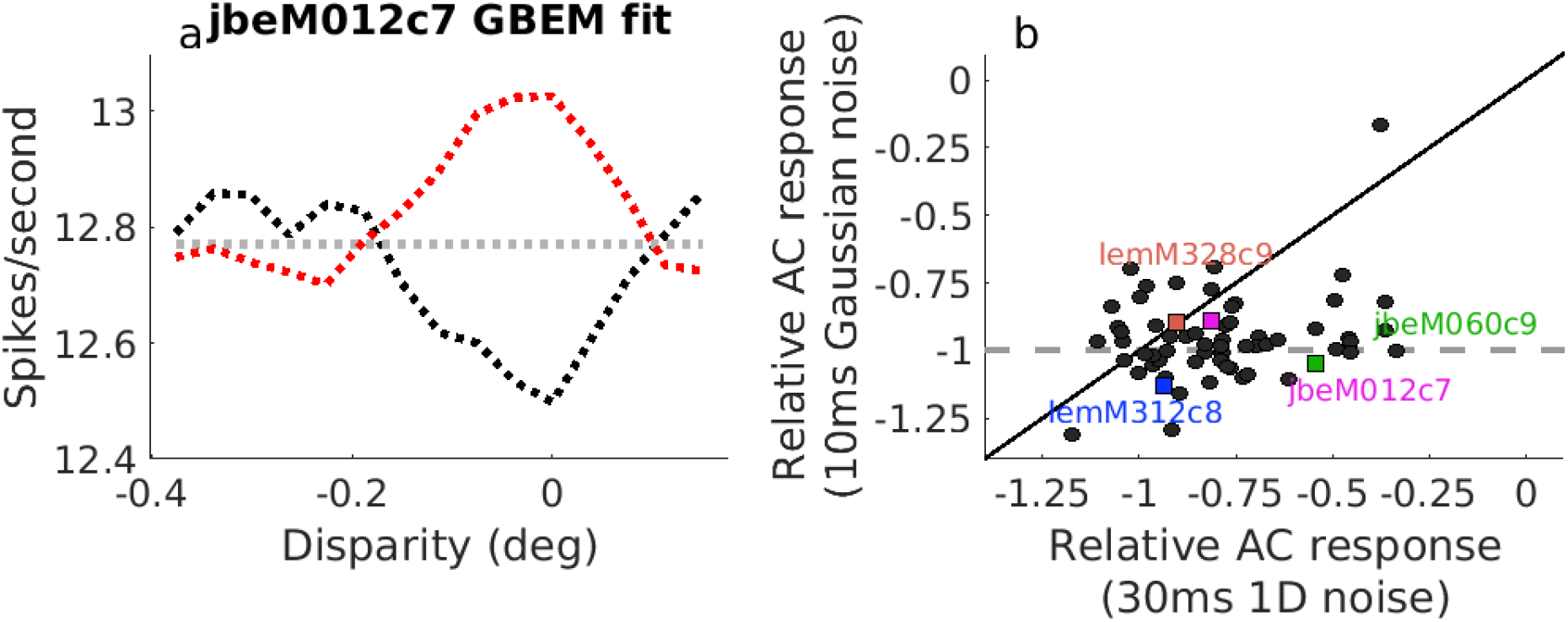
The predicted disparity tuning curve of cell jbeM012c7 in response to binocularly uncorrelated Gaussian noise (using forward correlation as in Tanabe et al), based on the GBEM fit. The tuning curve of the cell and its GBEM fit to 30ms 1D noise stereograms are shown in Figure 3c and d, respectively, where strong attenuation is present. Note that this attenuation is not present in model responses to binocularly uncorrelated Gaussian noise. b) The population summary for all GBEM units. The relative anticorrelated response to 1D noise is shown on the horizontal axis, and the relative anticorrelated response to Gaussian noise is shown on the vertical axis.

We can summarise this across the population by plotting the relative anticorrelated response to the 1D RLS stimulus against the relative anticorrelated response to the independent Gaussian noise stimulus (Figure 7). Just as in Tanabe & Cumming (2011), our model units cluster around −1 (no attenuation), and at the population level is not significantly different from −1 (t(94) = 1.53, p = 0.13). Thus, our model units readily capture both the attenuated anticorrelated response/correlated amplification exhibited by real cells to 1D RLS stimuli with binocular disparity, and also the absence of that attenuation in binocularly independent Gaussian noise.

## Discussion

Linear-nonlinear (LN) cascade models have been very successful in describing many properties of cortical neurons [Hubel and Wiesel, 1962, Ohzawa et al., 1990, Cumming and Parker, 1997, Ringach et al., 1997, Sharpee et al., 2011, Ostojic and Brunel, 2011]. Although they rarely predict spike rates perfectly, they can quantitatively describe the shape of tuning curves for several stimulus parameters [Hubel and Wiesel, 1962, Adelson and Bergen, 1985, Ohzawa et al., 1990, Cumming and Parker, 1997]. Thus existing data are compatible with a view in which LN-cascades generate cortical stimulus selectivity, while other nonlinearities only regulate the response gain. Invoking these nonlinearities has been reasonable, since the cascade models are usually fit using responses to stimuli that differ from the tests. Here we provide a new test of LN cascade models by fitting models to activity of V1 neurons in response to random noise patterns with binocular disparity, and then examining the mean responses of both model and cell across many different patterns of a given disparity.

Accounting for neuronal responses to disparity is made challenging by the fact that while V1 neurons modulate their activity with the disparity of anticorrelated patterns, this modulation is typically weaker than that observed in response to correlated patterns [Cumming and Parker, 1997]. A number of schemes have been proposed that might explain this [Nieder and Wagner, 2000, Read et al., 2002, Tanabe and Cumming, 2008, Tanabe et al., 2011, Tanabe et al., 2011, Samonds et al., 2013, Henriksen et al., 2016a, Burge and Geisler, 2014, Goncalves and Welchman, 2017], but it is unclear which, if any, of these model frameworks can account for neuronal activity in V1.

In this paper, we made use of developments in optimisation routines which allowed us to fit the components of generalised binocular energy model units to spiking data from V1 [McFarland et al., 2013]. The data was generally well-described by the model units, confirming that this class of models can serve as a first approximation. However, even the best-fitting models underestimated the magnitude of the correlated response at the preferred disparity of the cell. It is important to note that this is not simply a failure of the fitting procedure, such as a failure to find the global maximum likelihood. Although it is possible to create disparity tuning curves by hand that better capture the cell’s tuning curves, the GBEM was not optimised for fitting the actual tuning curves themselves. Rather, the GBEM finds a model which can best map image sequences to firing rates, and in doing so captures temporal and monocular dynamics, as well as binocular interactions. A model with a sufficiently wide array of LN filters must necessarily produce a good description of the responses to which it is fit, including the disparity tuning, since the architecture can approximate any function [Hornik, 1991]. For an LN cascade model to provide a realistic account of cortical neurons, it must both use a plausible number of subunits and generate accurate predictions to images which were not used in fitting the model.

We observed that the tendency for the GBEM to underestimate the correlated response was related to how much response attenuation the cell exhibited: cells that show relatively weaker modulation to anticorrelated stimuli were fit more poorly by our model framework. However, this was specific to the correlated responses. In other words, when cells had very large correlated responses relative to anticorrelated responses, the model failed to capture the correlated response to a larger degree than the anticorrelated response. This strongly suggests that disparity-selective V1 cells have a specialised mechanism for strengthening the response to correlated stimuli at the preferred disparity, which likely does not originate from a standard linear-nonlinear cascade. We call this mechanism correlated amplification.

Previous work has expanded on the BEM to account for the relatively weaker anticorrelated responses in four separate ways. The first and simplest is by simply appending an output nonlinearity to the BEM. This is by far the most common method for generating anticorrelated attenuation (e.g. [Nieder and Wagner, 2000, Read et al., 2002, Henriksen et al., 2016a]), but there has been little evidence showing that this is the actual explanation. Notably, a simple output exponent can be easily approximated by the thresholded square nonlinearity on the subunits in the GBEM. Therefore, if an output nonlinearity was the correct explanation, our modelling approach would have revealed it.

Anticorrelated attenuation in cells with odd-symmetric disparity tuning (with a peak at one disparity but a trough at another disparity) is a particular challenge [Read et al., 2002]. An output nonlinearity cannot account for the anticorrelated attenuation in these cells since the nonlinearity would affect the correlated and anticorrelated responses equally (and produce no attenuation). The only published solution is to combine two or more even-symmetric subunits with different preferred disparities [Haefner and Cumming, 2008, Read et al., 2002]. If each subunit shows attenuation (because of an output nonlinearity, or monocular thresholding), then this is present in the final response. If this two-subunit model were the correct description of odd-symmetric disparity tuning, then our fitting procedure should have recovered subunits with offset, even-symmetric disparity tuning. Instead, neurons with odd-symmetric tuning curves tended to have subunits which themselves had odd-symmetric tuning (e.g. lemM322c1 in Figure 1), and thus little anticorrelated attenuation (e.g. lemM328c9 and lemM312c8 in Figure 3). This finding argues against the two-subunit model of odd-symmetric disparity tuning.

Thus, our data show that a GBEM architecture can only partially account for observed responses to anticorrelation. A different approach was described by Samonds et al. (2013), who used a recurrent network made up binocular energy model-like units. Recurrent models are interesting because they can capture a range of complex phenomena known to operate in cortex. For example, feedforward models such as the GBEM are not ideal for modelling complex temporal dynamics. In contrast, Samonds et al. (2013) used their recurrent binocular network to successfully model the sharpening of disparity tuning over time [Samonds et al., 2009, Samonds et al., 2013]. Recurrent models are particularly promising given the recent success of recurrent neural networks in matching or exceeding human performance on a range of of complex visual tasks, such as image captioning [Vinyals et al., 2014] and object recognition [Liang and Hu, 2015]. Cells with recurrent connections remain a strong potential candidate for accounting for disparity tuning in real cells. However, this explanation has not yet been tested in real V1 neurons by fitting a complete recurrent model to individual cell responses.

One possibility that has not been explored is incorporating processes such as contrast normalisation separately for each subunit into LN cascades. It has been known for a long time that real cells have strong (largely monocular) contrast gain control [Albrecht and Hamilton, 1982, Simoncelli and Heeger, 1998, Truchard et al., 2000, Carandini and Heeger, 2012]. In other words, a large increment in contrast creates a relatively small increment in neuronal response, which is unlike what happens in models like the GBEM (where the response can change several orders of magnitude due to the quadratic nonlinearities). A second possibility is that more complex nonlinearities may be able to explain the correlated amplification process. Nonlinearities which act on multiple subunits have so far received little attention. For example, a cell whose final spiking nonlinearity is a thresholded AND gate, i.e.

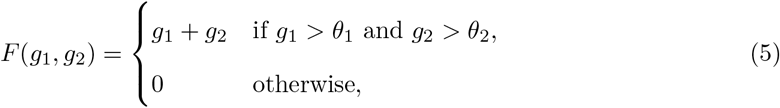

could potentially display the correlated amplification effect given appropriate subunits [Archie and Mel, 2000]. Similar nonlinearities have been explored in relation to contrast normalisation, but little attention has been given to these types of computations in describing selectivity for stimulus features.

A final unexplored possibility is that adding layers to the GBEM framework will allow it to learn more complex models which still generalise well to unseen data. Indeed, one of the remarkable features of deep convolutional neural nets is their ability to learn complex, nonlinear features from unstructured data [LeCun et al., 2015]. It is well-known that given enough subunits, even models such as the GBEM can in principle approximate any arbitrary function [Hornik, 1991]. However, if the model is not a good approximation of the underlying generative process, then the learned model parameters will fail to generalise to new, unseen data. For a deep generalised binocular energy model framework the question therefore becomes whether such a model can 1) provide a compact account of disparity-selectivity in cortex, and 2) help illuminate the mechanisms which gives rise to interesting properties of real cells.

Thus, while our results demonstrate that a simple LN cascade cannot provide a complete account of cortical stimulus selectivity even in V1, they also point the way to tractable related architectures that may provide a sucessful account, and a model system in which the model can be fit and tested using responses to a single set of stimuli. In several cases these preserve the initial linear stage (as proposed by Hubel and Wiesel for simple cells), but required a more complex interaction across subunits than simple linear summation. This interaction could be a unique contribution of cortex in generating stimulus selectivity.

## Footnotes

1 In theory, a cell whose anticorrelated response is modulated orthogonally to the correlated response would have a relative anticorrelated response of 0; in practice, the latter is rare and we observed no such cases in the current study.

